# Two sequence- and two structure-based ML models have learned different aspects of protein biochemistry

**DOI:** 10.1101/2023.03.20.533508

**Authors:** Anastasiya V. Kulikova, Daniel J. Diaz, Tianlong Chen, T. Jeffrey Cole, Andrew D. Ellington, Claus O. Wilke

## Abstract

Deep learning models are seeing increased use as methods to predict mutational effects or allowed mutations in proteins. The models commonly used for these purposes include large language models (LLMs) and 3D Convolutional Neural Networks (CNNs). These two model types have very different architectures and are commonly trained on different representations of proteins. LLMs make use of the transformer architecture and are trained purely on protein sequences whereas 3D CNNs are trained on voxelized representations of local protein structure. While comparable overall prediction accuracies have been reported for both types of models, it is not known to what extent these models make comparable specific predictions and/or generalize protein biochemistry in similar ways. Here, we perform a systematic comparison of two LLMs and two structure-based models (CNNs) and show that the different model types have distinct strengths and weaknesses. The overall prediction accuracies are largely uncorrelated between the sequence- and structure-based models. Overall, the two structure-based models are better at predicting buried aliphatic and hydrophobic residues whereas the two LLMs are better at predicting solvent-exposed polar and charged amino acids. Finally, we find that a combined model that takes the individual model predictions as input can leverage these individual model strengths and results in significantly improved overall prediction accuracy.

## Introduction

Machine learning models such as neural networks are increasingly being used to computationally explore protein variants suitable for protein engineering and to predict their effects on protein structure and function^1–6^. In fact, several deep neural network models, including large language models (LLMs) and convolutional neural networks (CNNs), have helped with the engineering of proteins with enhanced function/stability^7–10^. Interestingly, the deep learning models employed to date differ substantially in their architectures and, more importantly, the type of input data they are trained on. While LLMs are typically trained on large amounts of sequence data, CNNs and other model architectures have also been trained on input consisting of protein structures^11–13^, and models trained on both types of input data have produced meaningful insight on protein variants.

Sequence data is significantly more abundant than structure data, so it would be convenient if models could be trained purely on sequence data. And in fact, sequence-based models, and specifically protein large language models (protein LLMs) using transformer architectures, have been successfully employed in a number of contexts, including predicting variant effects and protein fitness^5,14,15^, predicting post-translational modifications and biophysical attributes^4^, predicting protein structure^4,16,17^, and even predicting entire protein-protein complexes^18^. At the same time, CNNs trained on structure data have been successful in enhancing enzyme function activity by suggesting activity enhancing protein variants^9,10^. Because sequence- and structure-based models learn from different protein representations, it is not clear whether they have inherent differences or instead make substantially equivalent predictions. Additionally, because the different model architectures have typically not been applied head-to-head to the same problem, excellent performance of one architecture in one problem area does not imply that a different architecture might not perform similarly well. Further progress on this question requires a direct comparison of different model architectures on the same problem.

Here, we compare two existing sequence-based and two existing structure-based models head-to-head on the task they were originally trained for, the prediction of a masked residue. We use two different sequence-based LLMs, protBERT^4^ and ESM1b^5^, and we compare them to two structure-based 3D CNN models, which we refer to here as CNN^19^ and RESNET^12^. We find that all these models generally predict the masked residue with similar accuracy. However, because the models differ in how they “see” protein biochemistry and what they take as input data, they tend to make distinct predictions that reflect different aspects of the underlying protein biochemistry. Further, we show that we can improve model performance by combining the models into a single, joint model. The combined model can compensate for weaknesses of the individual models by learning how to integrate the various model predictions and preferentially relying on the specific predictions that it estimates to be the most accurate for a given site in a given protein structure.

## Results

We compared the performance of four self-supervised deep neural network models on their original training task: predicting masked residues in proteins. The four models included two LLMs, ESM1-b^5^ and protBERT^4^, and two structure-based models, a 3D CNN^19^ and a Residual Network (RESNET)^12^. Our first question was whether any one of the models was consistently outperforming the others. We generated predictions for every residue in a test set of 147 protein structures. We found that the average accuracy across the 147 structures was 60.74%, 64.4%, 64.8% and 68.3%, respectively, for the ESM1b, CNN, RESNET and protBERT models (Fig. 1a). Here, we defined accuracy as the fraction of correct predictions across all residues in a single protein. A prediction is classified as “correct” if the predicted amino acid is identical to the amino acid that was originally masked (i.e., the wildtype amino acid), and the predicted amino acid is defined as the specific amino acid (out of all twenty possible) that is assigned the highest probability score by the model. Importantly, we found that prediction accuracy varied widely across protein structures, in particular for the LLMs, which yielded accuracies as low as 0.2 for some structures and in excess of 0.9 for other structures. Accuracies were more consistent for the CNN and RESNET models, ranging from approximately 0.5 to 0.8. We believe these observations reflect the well-known bias/variance dilemma in machine learning^20^: Convolution layers innately possess inductive bias for spatial data whereas sequence-based transformer architectures, although more powerful, lack this bias and therefore their inferences tend to display higher variance.

**Figure 1.**
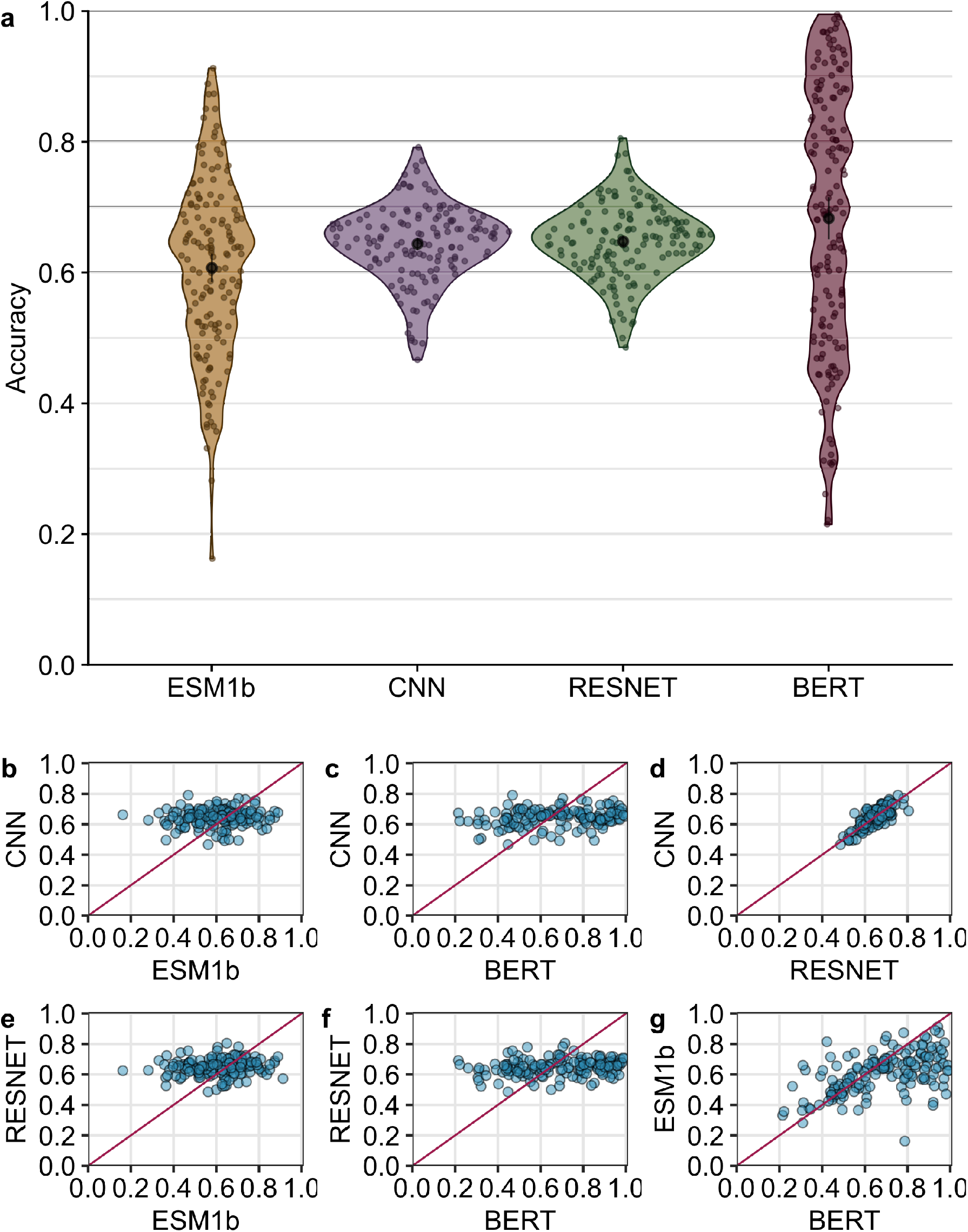
Prediction accuracy compared across models. (a) Average accuracy per protein for each model. The average accuracy is 60.74%, 64.4%, 64.8%, and 68.3%, respectively, for the ESM1b, CNN, RESNET, and BERT models. Average accuracies are highlighted by the black point within each violin. (b) Correlation between CNN and ESM1b accuracy (*r* = 0.09, *p* = 0.28). (c) Correlation between CNN and BERT accuracy (*r* = 0.22, *p* = 0.008). (d) Correlation between CNN and RESNET accuracy (*r* = 0.77, *p <* 10^−10^). (e) Correlation between RESNET and ESM1b accuracy (*r* = 0.12, *p* = 0.16). (f) Correlation between RESNET and BERT accuracy (*r* = 0.20, *p* = 0.016). (g) Correlation between ESM1b and BERT accuracy (*r* = 0.52, *p* = 10^−10^). Each point represents a single protein.

We next asked whether the four models had comparable accuracies for the same protein structures. In other words, if one model generated either high or low accuracy for a given protein, did the other models do the same, or did they behave differently? We found that the structure-based models and the LLMs displayed markedly different behavior. The prediction accuracy of neither of the transformer models was correlated with the accuracy of the CNN or RESNET models (Fig. 1b, c, e, f). By contrast, there was a moderate correlation between the predictions of the two LLMs (Fig. 1g) and a strong correlation between the two structure-based models (Fig. 1d). These findings imply that the two LLMs, which share their input data type and general aspects of their architecture (both are transformer-based language models), display similarities in their predictions but behave by no means identically. And the structure-based models make entirely different, uncorrelated predictions to the LLMs. We would like to emphasize that the two structure-based models displayed a strong correlation even though they were trained on different datasets, using different microenvironment sampling procedures, and they also had substantially different architectures. The main similarities between these two models are that they use structure-based data as input, convolution layers rather than attention for processing, and training data curated with at most 50% sequence similarity.

Because predictions between model types were uncorrelated, we next asked whether we could combine outputs from the four models for improved prediction accuracy overall. Initially, we tried very simple ensemble methods, such as averaging the predictions among models or taking the highest probability among individual models as the final prediction. Neither method yielded meaningful improvements in overall prediction accuracy over the individual models, and therefore we concluded that a more sophisticated ensembling approach was needed.

We trained a simple fully-connected neural network model to produce a combined prediction from the four individual models. The combined model takes in 80 probabilities (20 per input model), passes them into two intermediate dense layers, and ultimately outputs a set of 20 probabilities, where each node represents the probability of one of 20 amino acids (Fig. 2). We trained this model (Supplementary Figs. S1 and S2) using a training dataset consisting of 3,209 proteins with a sequence similarity of at most 80% to any of the 147 proteins in the test set or the training sets of the individual models. After training, we generated predictions on a test set of 147 proteins and assessed prediction accuracy for each protein. We found that the combined model outperformed all four individual neural networks with an average accuracy of 82%, an improvement of over 10 percentage points from the most accurate individual model (Fig. 3).

**Figure 2.**
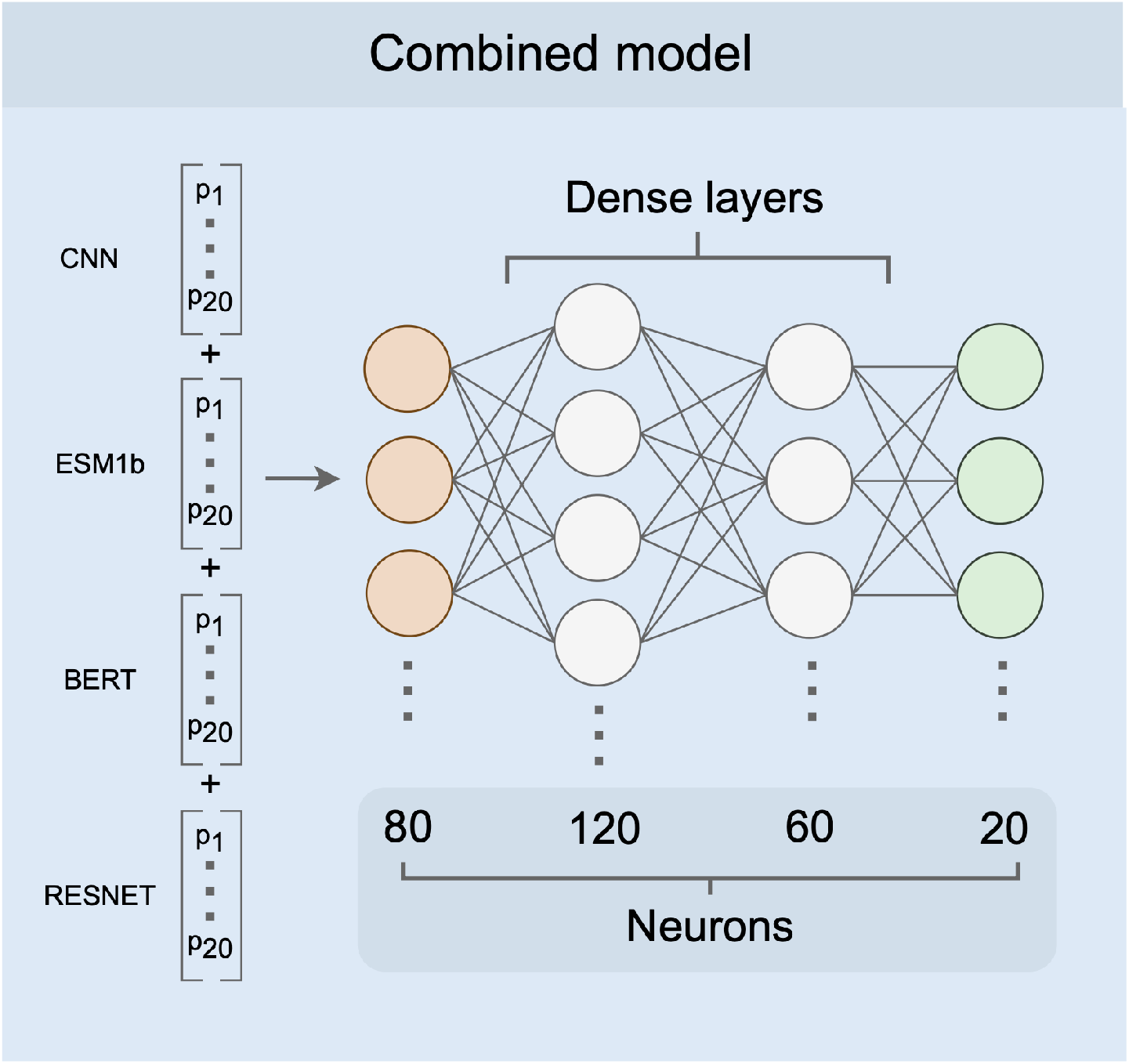
Combined model architecture. As input, the combined model receives a conjoined input vector of length 80 (orange). There are two hidden layers with 120 and 60 nodes, respectively (white). The output is a vector of 20 probabilities, one for each of the amino acids (green).

**Figure 3.**
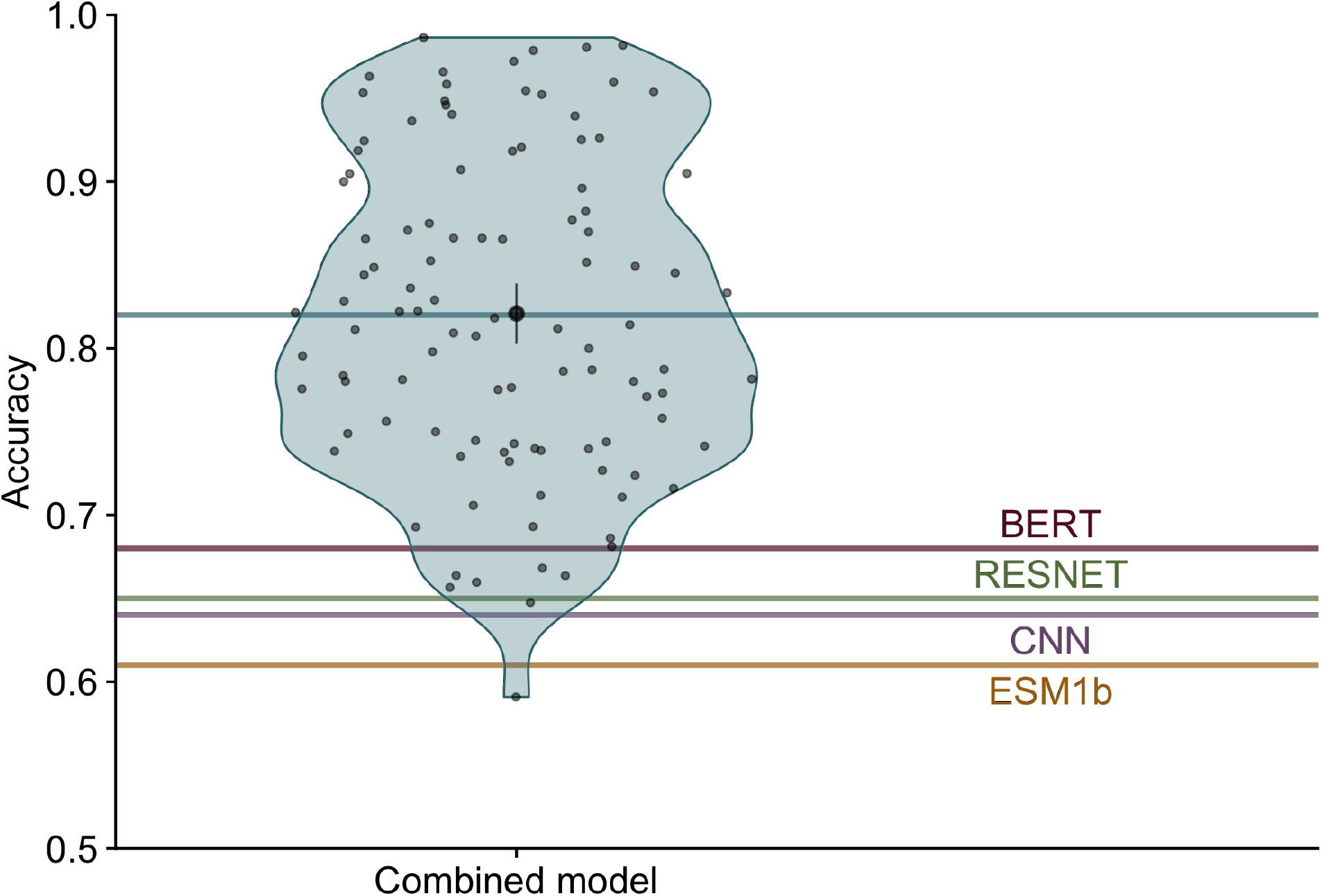
Combined model accuracy. Each blue point within the violin shows the prediction accuracy for a single protein. The mean accuracy across all proteins, indicated by the blue line, was 82%. The black point and bars represent the mean and 95% confidence interval, respectively. Accuracies of individual models are indicated by red (BERT), green (RESNET), purple (CNN) and yellow (ESM1b) lines.

We next assessed how well the different models predict specific amino acid classes. We pooled all sites from all 147 test proteins and divided the sites by amino acid class (defined in Supplementary Table S1). We then calculated the accuracy within each amino acid class for all four individual models as well as the combined model. We found that overall, the combined model improved predictions for all classes and outperformed all individual models (Fig. 4). When looking at individual models, we see that the structure-based models outperformed the language models for aliphatic and unique (G and P) amino acids, while language models generally outperformed the structure-based models for polar and charged amino acids.

**Figure 4.**
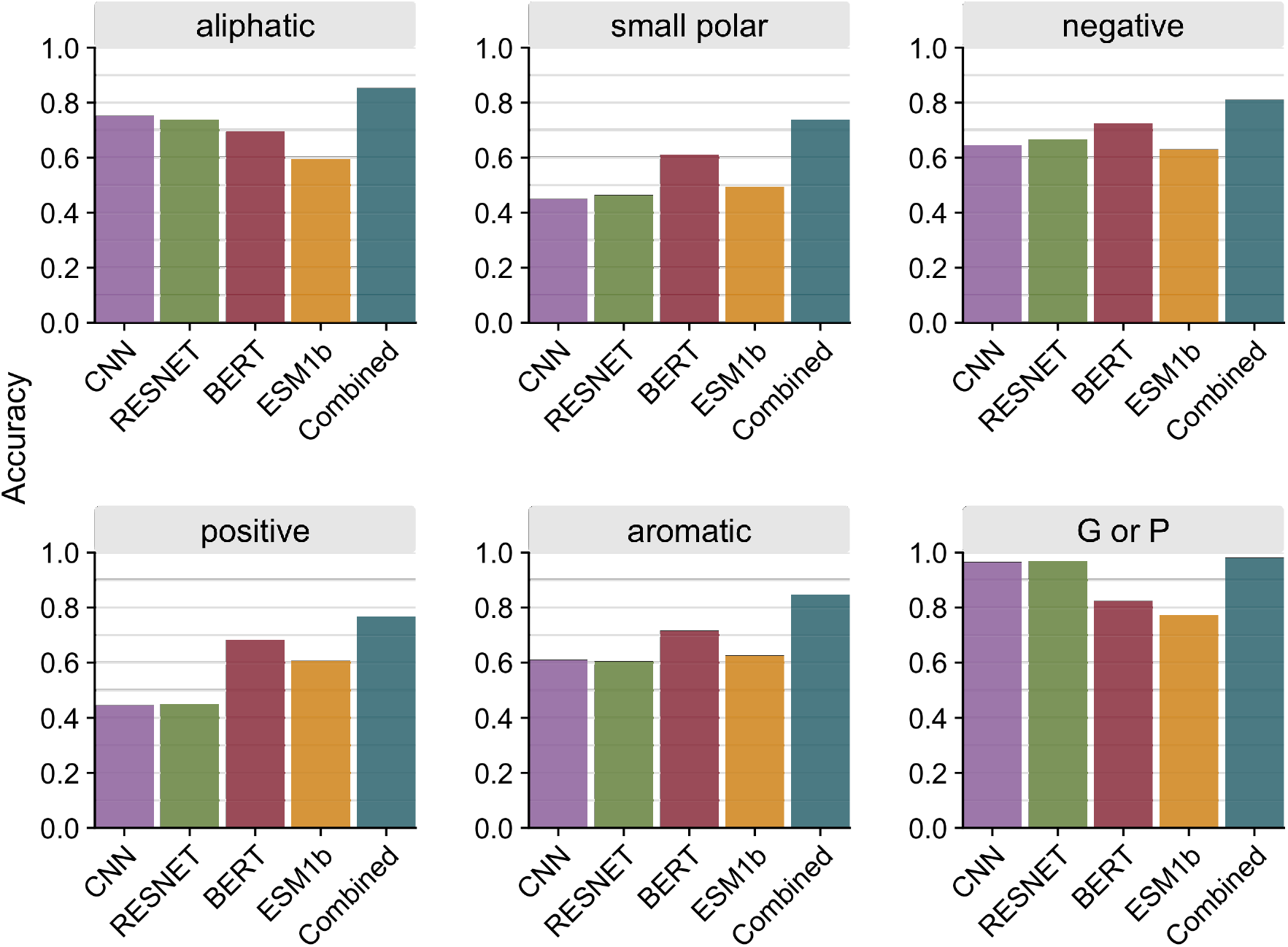
Prediction accuracy per amino acid class. Accuracy was calculated as an average across all sites of a specific amino acid class for each model. Sites were pooled together from all 147 proteins in the test dataset.

We then inspected how the accuracy of combined predictions relates to the individual predictions. To do so, we classified every individual prediction by whether the combined model agreed with all four individual models, only structure-based models, only sequence models, or none of the model. Unsurprisingly, when all models were unanimous, the prediction accuracy was very high, 96.9% (Fig. 5a). In other words, whenever the four individual networks individually made the same prediction, and that prediction also coincided with the combined network, then that prediction was likely correct. This scenario was also by far the most common, occurring 38.3% of the time (Fig. 5b). We saw the next highest accuracies whenever the combo model prediction agreed with at least one the transformers, closely followed by the case where a combination of structure and language models agreed with the combined model and the case when one or both structure-based models agreed (Fig. 5a). For these predictions, the combined model was correct between 70% to 80% of the time. On the other hand, unique predictions (where the combined model did not agree with any of the individual models) were the least likely correct, yet still better than random chance (5%) at an accuracy of ∼32.7%.

**Figure 5.**
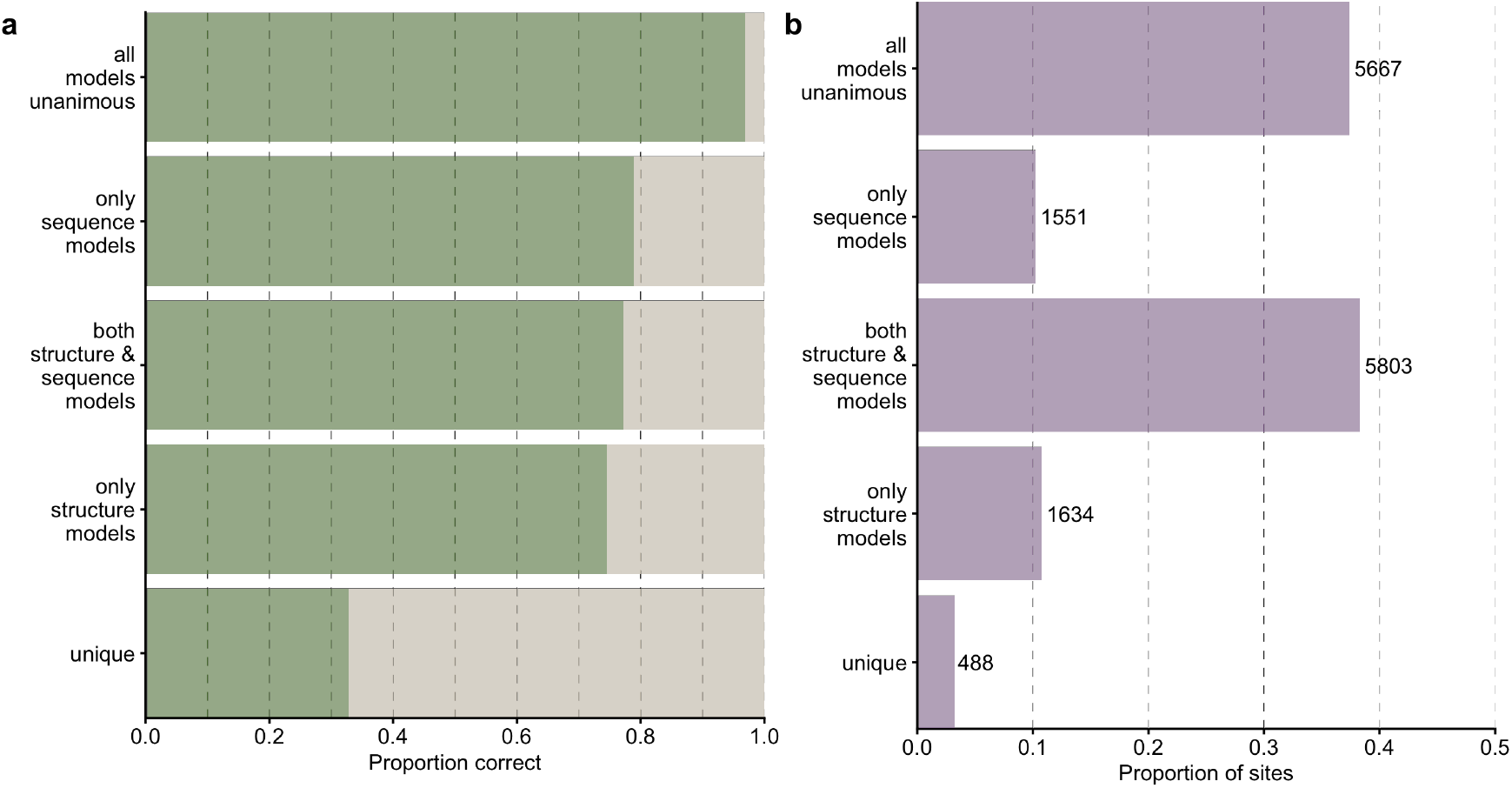
Comparison of combined model predictions with individual model predictions. (a) Proportion of correct predictions for sites at which the combined model prediction agrees with predictions of specific model combinations. (b) Proportion and number of sites corresponding to each scenario under part (a).

Thus, all possible combinations of individual model predictions were possible as predictions by the combined model, including the case where the combined model disagreed with all individual models. This observation suggests that the combined model learns how to interpret the amino acid probability distribution for each individual model to properly aggregate the information into a more generalizable distribution rather than simply preferring one individual model or applying majority rule. We further confirmed the differences between LLMs and structure-based models by breaking down the distribution of correct predictions by individual amino acids (Supplementary Fig. S3). This analysis can be thought of as asking which individual model the combined model relied on to make specific predictions. We found that in cases where the combined model relied exclusively on the structure-based models’ predictions, the correctly predicted amino acids were more likely aliphatic, whereas in cases where the combined model relied exclusively on predictions from the LLMs, the correctly predicted amino acids tended to be charged (positive or negative) or polar. Somewhat surprisingly, the correct predictions made by the structure-based models displayed an amino acid distribution most similar to the overall distribution of amino acids within the proteins (Supplementary Fig. S4). The distribution of amino acids when all models were in agreement was somewhat similar (Supplementary Fig. S3). By contrast, the distribution of amino acids correctly predicted by one or both LLMs were markedly different (Supplementary Fig. S3). Finally, for correct unique predictions, where the combined model was correct and all individual models where incorrect, the shape of the distribution of amino acids was more similar to the distribution of correct LLM predictions than to the overall distribution of amino acids in the proteins (Supplementary Fig. S3).

Finally, we asked whether the solvent exposure of a site (i.e., whether it is on the surface or in the core of the protein) contributed to differences in predictions between the models. We did this by correlating the Relative Solvent Accessibility (RSA) of each site in our dataset with the model confidence for that site. Model confidence is defined as the probability with which the top scoring amino acid is predicted at a site, and it correlates well with model accuracy (Supplementary Fig. S5 and prior work^19^). Therefore, we used it here as an approximation of model accuracy, which cannot be defined for individual sites (a site either is or is not predicted correctly). We found that all models had a tendency to have higher confidence in their predictions for buried residues (RSA near zero) than for exposed residues (RSA of 0.2 or larger) (Fig. 6). However, on average, performance of the RESNET and BERT model was more uniform across the RSA range (and in particular for the RESNET model) than it was for the CNN and ESM1b models. Both the CNN and ESM1b model made a large number of low-confidence predictions above RSA of 0.2. In the combined model, we saw prediction confidence markedly increased across all RSA values. The network did not have a strong bias towards either buried or exposed residues and instead made predictions with comparable confidence across the entire spectrum of RSA values.

**Figure 6.**
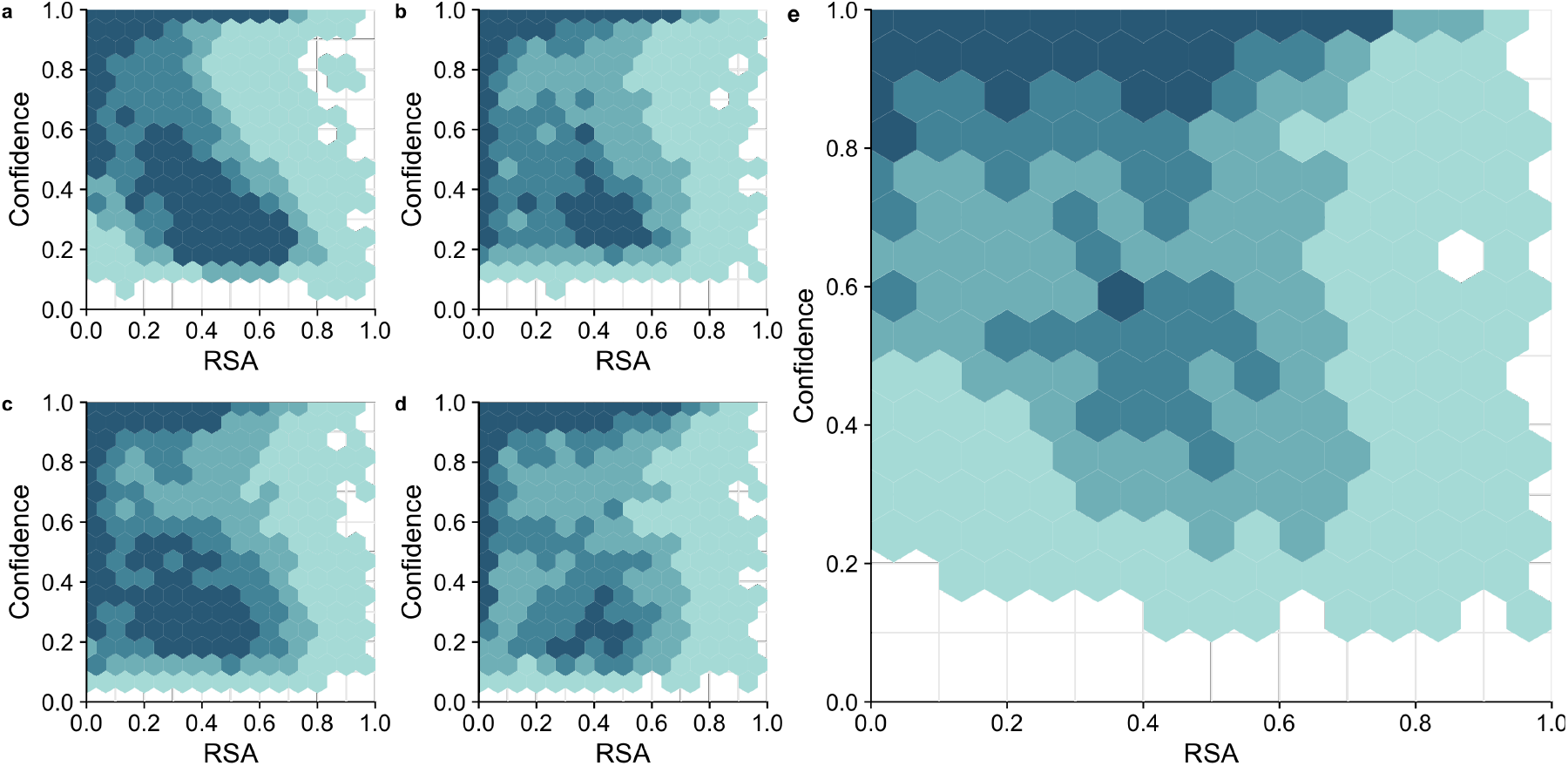
Model confidence as a function of Relative Solvent Accessibility (RSA). (a) CNN (b) RESNET (c) ESM1b (d) BERT (e) Combined model. Counts are binned into the following four groups: 1–25 predictions/site (lightest color), 26–50 predictions/site, 51–75 prediction/site, and over 75 predictions/site (darkest color).

## Discussion

We have compared the performance of two existing structure- and two existing sequence-based neural network models on the task of predicting masked residues in proteins, the task that all these models were originally trained on. We have found that the two different model types vary widely in their performance on specific proteins, even if their average accuracies are similar. Predictions by structure-based models are not correlated or at most weakly correlated to predictions by sequence-based models. The structure-based models perform best at predicting aliphatic core residues, whereas the sequence-based models provide more accurate predictions for polar, solvent-exposed sites. We have further constructed a combined model that uses the predictions of the individual models as input and turns them into an ensemble prediction. This combined model has achieved significantly higher prediction accuracy than the individual models, and importantly, can leverage the individual models for their specific strengths. The combined model seems to be able to select the individual model predictions that are most suitable for specific sites.

Sequence-based language transformer models and structure-based convolutional models differ widely in their training data, their internal structures, and their strengths and weaknesses. The transformer architecture has grown popular in the fields of natural language processing and image recognition^21,22^. More recently, the self-supervised technique masked-language modeling (MLM) used to train LLMs has been applied to predict masked residues in protein sequences^3–6^. Such LLMs can capture interactions between amino acids that are far apart in the sequence but spatially close in the structure, relying solely on input sequence for inference, without the need for additional features or annotations^17^. By contrast, 3D CNNs are fundamentally different from LLMs as they take the local protein structure rather than the entire amino-acid sequences as input^8–10,12,19,23^. The input consists of a voxelized box built around a focal residue which is removed before training or inference. The advantage of this structural data as model input is that it directly provides information on physical contacts within a protein, which are most critical for accurate residue prediction^8,19^.

Because LLMs do not use any structural information as input, we might expect them to perform consistently worse than 3D CNNs at predicting masked residues. However, we have seen that this is not the case for the four models considered here; both model types predicted residues with comparable overall accuracies. At the same time, there were clear differences between model types. The structure-based models tended to perform better at predicting aliphatic or hydrophobic residues, whereas the LLMs had an edge for polar or charged amino acids. These differences may be due in part to the fact that the latter amino acids are more common on the surface of the protein, where the voxelized input boxes to the structure-based models are partially empty and thus provide less data to base inference on.

The consequence of the differing strengths and weaknesses of the two model architectures is that the different models can be combined for improved overall performance. The network output for all four models consists of a distribution of 20 probabilities, one for each of the amino acids. This distribution of probabilities can also be viewed as an embedding for a biochemical environment^24–26^. Considering that the distributions of probabilities can differ widely even if the same amino acid is predicted to be the most likely, we expect that these distributions carry meaningful information about the microenvironment surrounding the focal residue in the folded protein. Moreover, we would expect similar microenvironments to have similar distributions^27^. Consequently, the combined model can use the probability distributions of the four input models to infer the likely microenvironment of the focal residue and then infer which amino acid to place at this location. This is not a simple majority rule among the models but a non-trivial inference task, as can be seen from the fact that the model on occasion makes unique predictions distinct from any of the input models.

One challenge we encountered with LLMs is achieving a clean separation between training and test datasets. Because the LLMs require very large training datasets, they are frequently trained on virtually any unique protein sequence available. To put into perspective, ESM-1b (650M parameters) and ProTrans (450M parameters) were trained with ∼27M and *>* 80M protein sequences with 50% and 100% sequence similarity from UniRef^28^, respectively, whereas the structure-based models we used here (both with 12M parameters) were trained with ∼1.6M and ∼2.3M microenvironments sampled from ∼17K and ∼23K sequences with a 50% sequence similarity. The large training sets for the language models raise the possibility that these models overfit to their training datasets to the point of memorization. With enough training, any model will eventually memorize the training set and be unable to generalize to unseen data^29,30^. We saw evidence of potential overtraining when we used more recent models of protBERT and ESM1-b, as well as ESM1-v^6^. For these models, we obtained accuracies averaging at ∼95% across proteins (Supplementary Figs. S6 and S7). However, since these models had trained on datasets containing most proteins in the UniRef, we could not guarantee that our test sequences were not in the training data. In fact, we are quite certain they were. Here, primarily for this reason, we used older verisons of protBERT and ESM1-b because they were trained on smaller datasets. Ideally, all models would need to be retrained on a clean and consistent dataset for a better comparison.

In summary, our results show that two different neural network types, due to differences in how they learn protein features, can make unique contributions to the same residue prediction task. At the same time, predictions that match between a sequence-based and a structure-based model are most likely to be correct, implying that although model predictions are not strongly correlated they do show some overlap. It remains an open question to what extent our results here generalize to other models with similar or differing architectures, and whether LLMs can in principle be trained to the point where they perform as well as or better than a structure-based model. One conceptual difficulty with this question is that LLMs use such large training sets and so many parameters that they in essence memorize the entire known protein universe. To what extent such models actually generalize protein biochemistry is difficult to assess.

## Methods

We used four pre-existing self-supervised deep learning models trained to predict masked residues: two sequence-based transformer models, ESM1-b^5^ and protBERT^4^, and two structure-based models, CNN^19^ and RESNET^12^. The two structure-based models have been trained on the same dataset, while the LLMs have been trained on separate and distinct training datasets. For all models, we defined model performance as the accuracy with which each model could predict masked amino acids in an array of different proteins.

### Testing Individual Models

We first assessed each model separately, using a test dataset we had previously used in studying the 3D CNN model^19^. Our test set is derived from the PSICOV dataset^31^, which consists of 150 well studied protein structures commonly used for covariation analyses. From this dataset, we removed three proteins for technical reasons, as described^19^. We verified that none of the remaining 147 proteins were in UniRef50, the training set of the ESM1-b model. Similarly, the CNN and RESNET training set had been chosen to not contain proteins with a sequence similarity of greater than 50% to any of the proteins in the test dataset^19^. Because protBERT has been trained on nearly every sequence in the UniRef RCSB databank^28^, we could not guarantee that our test sequences were not in the training data for that model.

LLM output was generated using the `berteome` library (version 0.1.6) available at: https://github.com/tijeco/berteome. The berteome library requires prior installation of the transformers package available at: https://pypi.org/project/transformers/. We used version 4.10.0 of the transformers package. The berteome library generates ESM1-b predictions from the pre-trained esm1b_t33_650M_UR50S model^5^ and protBERT predictions from the pre-trained Rostlab/prot_bert model^4^. CNN model output was generated using the network with input box size of 20Å^3^ as described^19^. The CNN model is available at https://github.com/akulikova64/CNN_protein_landscape and the RESNET model is available at https://github.com/danny305/MutComputeX.

### Generating a Combined Model

After assessing each individual model, we used the outputs of all four neural networks as training data for a combined model. The combined neural network model consisted of four layers: an input of 80 nodes (20 per model), two intermediate layers of 120 nodes and 60 nodes (“relu” activation function), respectively, and a final output layer of 20 nodes (“softmax” activation function). Each node in the output layer represents the probability of one of the 20 amino acids. All layers are fully connected. The model was implemented in python (v3.8.9) with the Tensorflow library (v2.9.2) using the keras API (v2.9.0)^32^.

To compile a training set for the combined model we first downloaded all PDB ID’s of proteins clustered into 80% homology groups from the RCSB website using the following link: https://cdn.rcsb.org/resources/sequence/clusters/clusters-by-entity-80.txt. To avoid reusing any proteins previously used to train the individual models, we removed any proteins that were part of the training sets of the 3D CNN (training set as described^19^), RESNET and ESM1-b (UniRef50), as well as the test set (PSICOV) and their homologs of 80% or greater similarity. We did not filter out proteins used to train the protBERT model as it was pre-trained on the UniRef100 dataset, containing most of the proteins in the RCSB database. Proteins longer than 1024 residues were also removed due to length restrictions of the ESM1-b model. Finally, we removed protein structures where we could not add hydrogen atoms or partial charges with PDB2PQR (v3.1.0)^33^. The resulting training set contained a total of 3209 protein structures.

To produce training data for the combined model, we generated CNN and RESNET predictions for each residue in our 3209 protein structure files. Similarly, the sequences corresponding to the 3209 protein structures were used to generate predictions from the protBERT and ESM-1b models. The combined network was trained on each protein/sequence position in this dataset for 150 epochs, on the goal of being able to predict the residue at each position in a protein. We used a fixed learning rate of 0.0001 using the Adam optimizer and the “categorical crossentropy” loss function. Finally, the fully trained combined network was tested against the same set of 147 proteins we had used initially to test individual models.

Relative solvent accessibility values were calculated using freeSASA software^34^ and applying the normalization constants as described^35^.

## Data availability

Final data analysis and figure production was performed in R^36^, making extensive use of the tidyverse family of packages^37^. The trained neural network model, analysis scripts, training set, and processed data are available on GitHub: https://github.com/akulikova64/BERT_CNN_comparison/

## Acknowledgements

This work was supported by grants from the Welch Foundation (F-1654), the Department of Defense – Defense Threat Reduction Agency (HDTRA12010011), and the National Institutes of Health (R01 AI148419). T.C. was supported by a fellowship from the NSF’s National AI Institute for Foundations of Machine Learning (2019844). We would like to thank Advanced Micro Devices, Inc. (AMD) for the donation of critical hardware and support resources from its HPC Fund that made this work possible. C.O.W. also acknowledges funding from the Jane and Roland Blumberg Centennial Professorship in Molecular Evolution and the Dwight W. and Blanche Faye Reeder Centennial Fellowship in Systematic and Evolutionary Biology at UT Austin.

## Author contributions statement

A.V.K., T.J.C., and C.O.W. conceived of the study. D.J.D., T.C., and T.J.C. contributed materials. A.V.K. performed all data analysis and wrote the initial manuscript draft. All authors edited and reviewed the final manuscript.

