## Supplementary figures and tables for "Two sequence- and two structure-based ML models have learned different aspects of protein biochemistry"

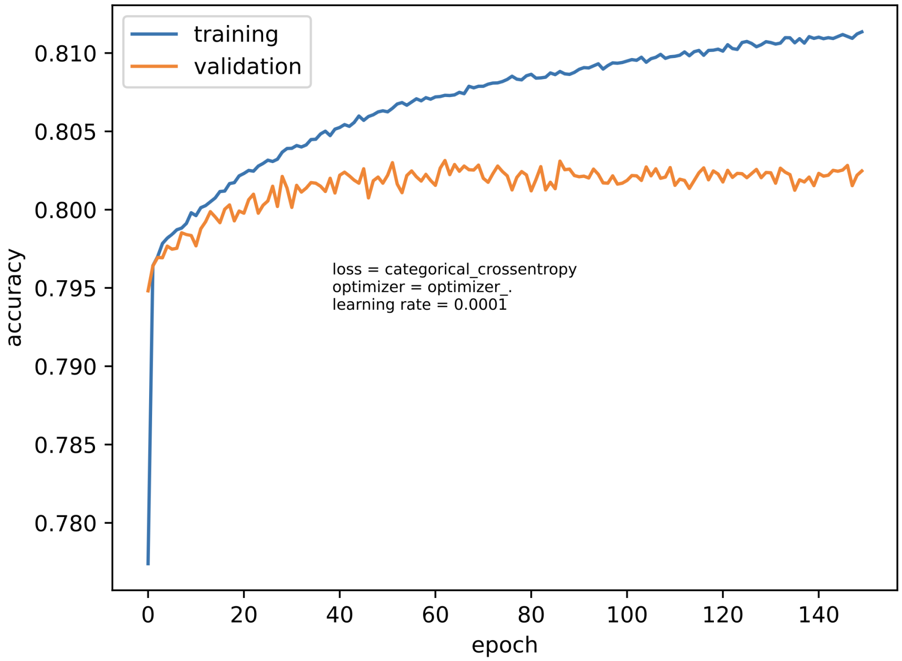


**Figure S1.** Combined model training and validation accuracy. The plot shows the training accuracy (blue) and the validation accuracy (orange) after each epoch for the combined model starting from epoch 1 to epoch 150. After epoch 1, training and validation accuracy were 77.74% and 79.48%, respectively. After epoch 150, validation accuracy reached 81.14% while training accuracy reached 80.25%.

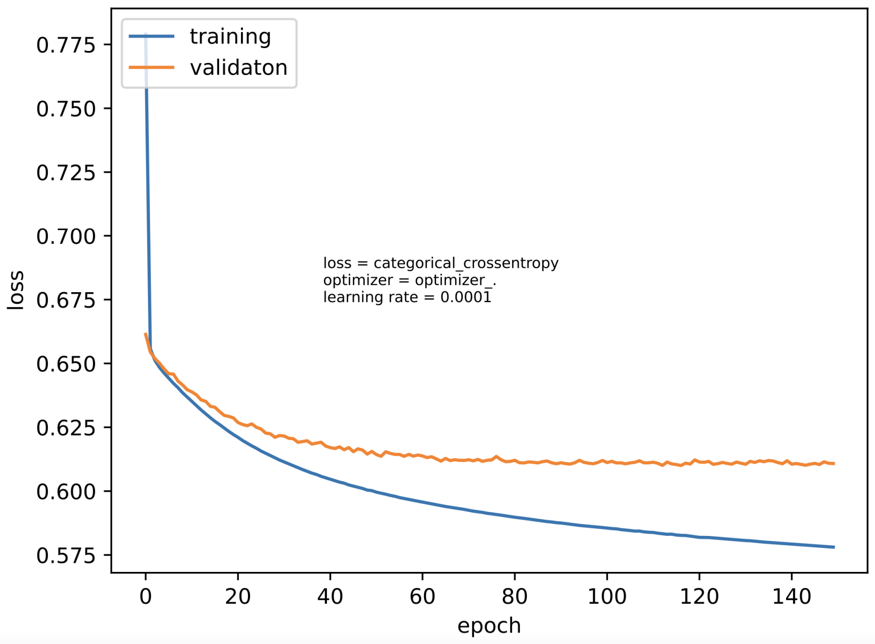


**Figure S2.** Combined model training and validation loss. The plot shows the training loss (blue) and the validation loss (orange) after each epoch for the combined model starting from epoch 1 to epoch 150. After epoch 1, validation loss was 0.6614, while training loss was 0.7790. After epoch 150, validation loss was 0.6108, while training loss was 0.5781.

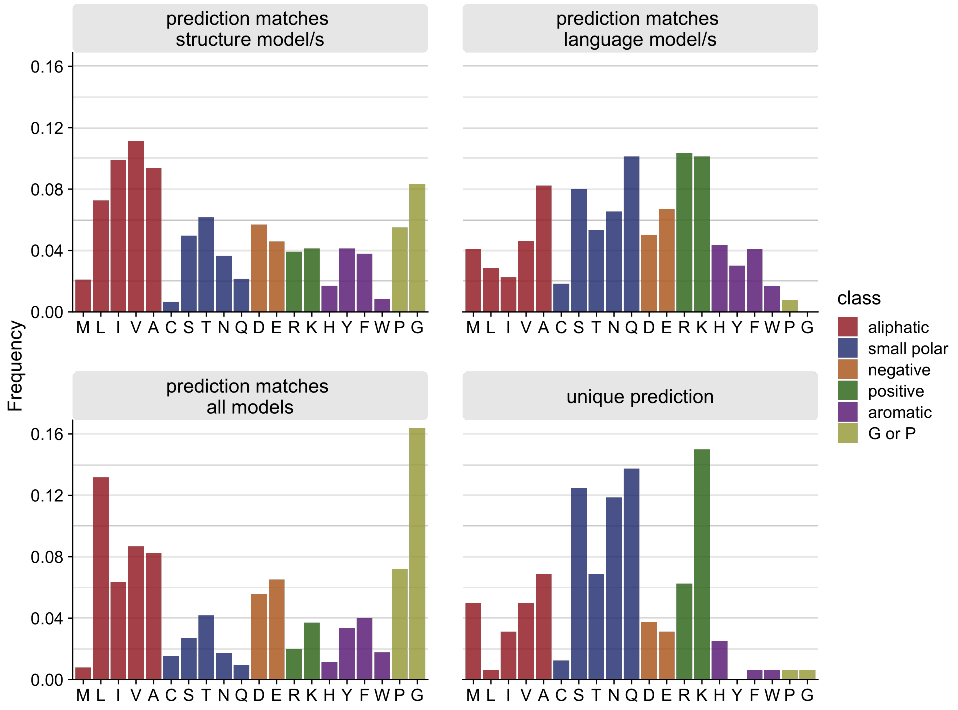


**Figure S3.** Relative frequencies of correctly predicted amino acids by the combined model, for cases where the prediction of the combined model matches (from left to right and top to bottom) the prediction of only one or both of the structure-based models, only one or both of the LLM models, all four individual models, or none of the individual models.

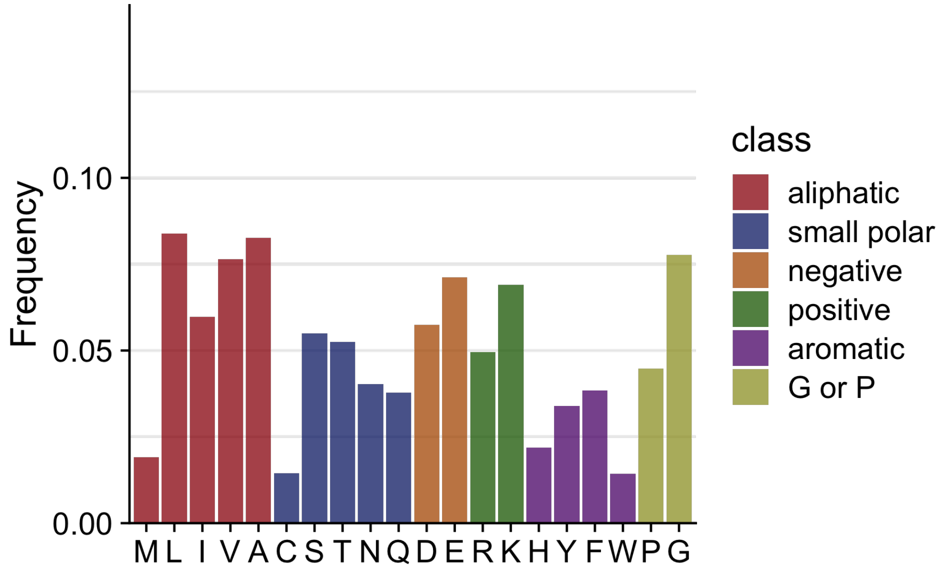


**Figure S4.** Distribution of wild type amino acids across 147 proteins from the PSICOV dataset.

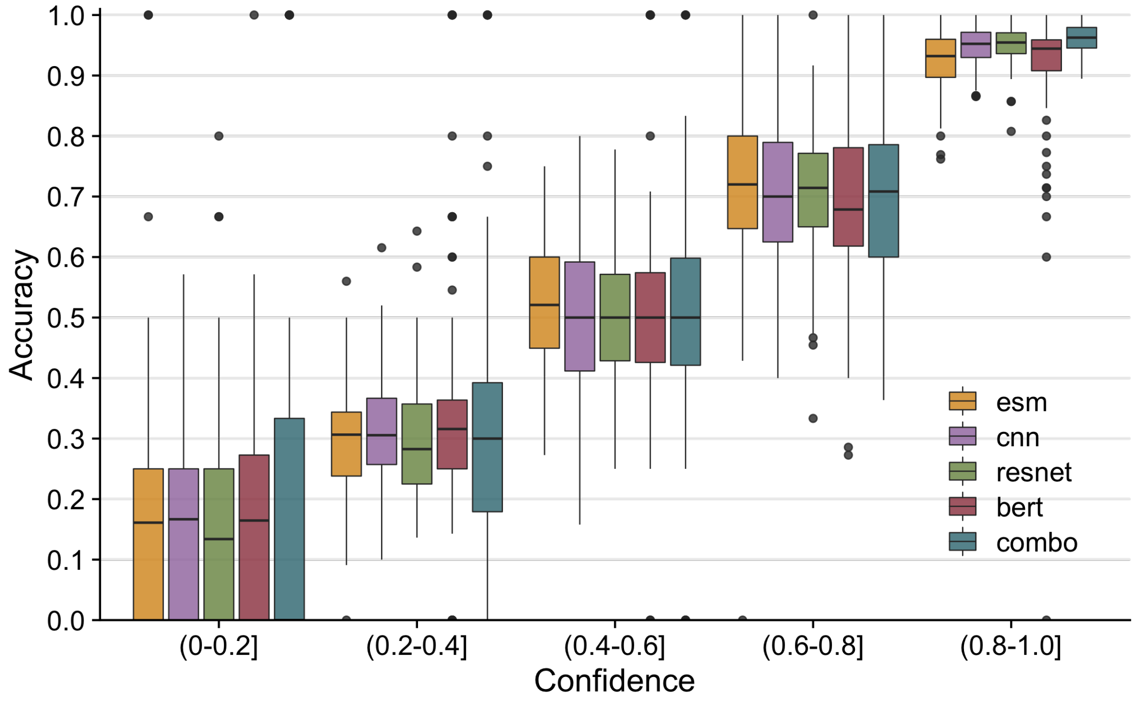


**Figure S5.** Prediction accuracy as a function of model confidence for all five models considered in this work.

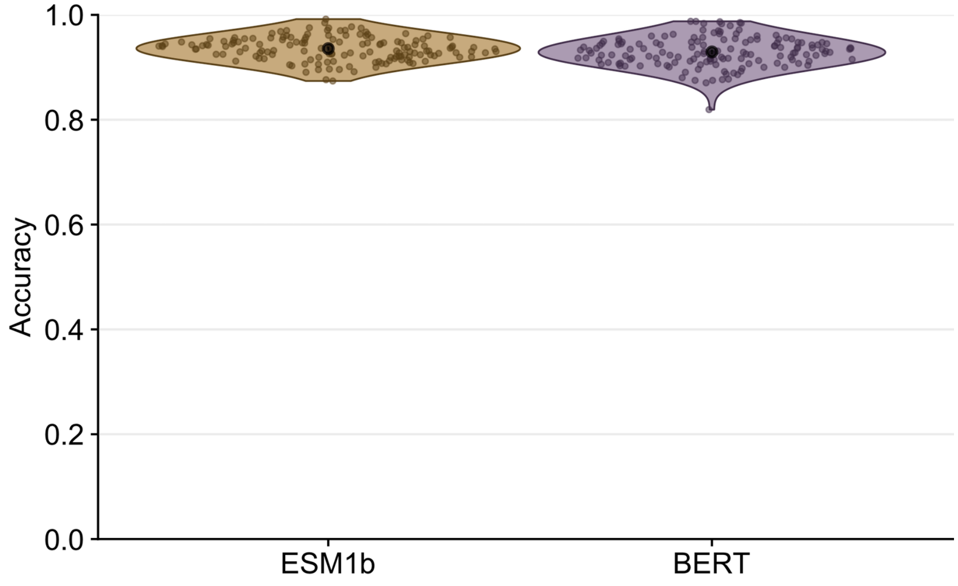


**Figure S6.** Average accuracy per protein of newer versions of transformer models. The average accuracy is ∼93% average accuracy for ESM-1b and ∼92% accuracy for the protBERT model. Each violin plot contains ∼150 data-points.

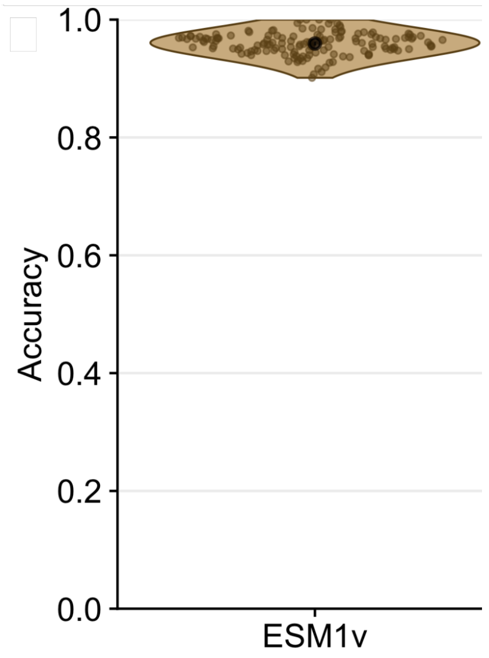


**Figure S7.** Accuracy of ESM1-v model. When the ESM1-v model (model: esm1v_t33_650M_UR90S_1) was tested on 147 proteins of the PSICOV dataset, the average accuracy for ESM1-v was ∼95%, as indicated by the black point in the center of the violin.

**Table S1.** Amino acids grouped by class. The following amino acid groups were used in Figure 4 and are mentioned throughout the paper.

| **class** | **included amino acids** |
|---|---|
| aliphatic | Methionine (M), Leucine (L), Inosine (I), Valine (V), Alanine (A) |
| small polar | Cysteine (C), Serine (S), Threonine (T), Asparagine (N), Glutamine (Q) |
| negative | Aspartic Acid (D), Glutamic acid (E) |
| positive | Arginine (R), Lysine (K) |
| aromatic | Histidine (H), Tyrosine (Y), Phenylalanine (F), Tryptophan (W) |
| unique | Proline (P), Glycine (G) |
